# Hollow condensates emerge from gelation-induced spinodal decomposition

**DOI:** 10.1101/2025.06.25.661497

**Authors:** Cheng Li, Lingyu Meng, Yongxin Tong, Jie Lin, Zhi Qi

**Affiliations:** Center for Quantitative Biology, Academy for Advanced Interdisciplinary Studies, Peking University, Beijing 100871, China; Peking-Tsinghua Center for Life Sciences, Academy for Advanced Interdisciplinary Studies, Peking University, Beijing 100871, China; The Integrated Science Program, Yuanpei College, Peking University, Beijing 100871, China; School of Physics, Peking University, Beijing 100871, China

**Author notes:** Equal contribution (Cheng Li, Lingyu Meng, Yongxin Tong).

## Abstract

Recent studies have identified diverse hollow biomolecular condensates, characterized by biomolecule-depleted interiors surrounded by biomolecule-rich shells. Although several formation mechanisms have been proposed, a general thermodynamic driving force remains elusive. Here, we investigate a well-defined system in which the human transcription factor p53 and non-specific double-stranded DNA (dsDNA) form biomolecule-rich condensates. Introduction of dsDNA containing p53-binding motifs induces a morphological transition to hollow structures, accompanied by a material state transition from liquid-like to gel-like. *In vitro* assays indicate that the formation of hollow condensates is driven by p21 DNA-induced localized gelation at the condensate periphery. Guided by these findings, we developed a three-component phase-field model that quantitatively recapitulates the formation of hollow condensates. Simulations show that peripheral gelation leads to gradual depletion of protein and Random DNA from the condensate core, triggering spinodal decomposition and lumen formation inside condensates. Together, these results offer mechanistic insights into multi-component hollow condensates.

## Introduction

Nucleic acids and proteins can assemble into biomolecular condensates through phase separation, a process that has been shown to play critical roles in various biological processes ^1^. Dysregulation of condensate formation is increasingly implicated in the pathogenesis of various human diseases, including neurodegenerative disorders and cancer ^2^.

In addition to forming biomolecule-rich condensates, biomolecular condensates can also assemble into hollow architectures, in which a biomolecule-depleted region is created within the biomolecule-rich condensate ^3^. For example, Banerjee and colleagues ^4^ demonstrated the formation of stable hollow condensates consisting of an arginine-rich disordered nucleoprotein, protamine (PRM), in combination with RNA. They proposed that protamine–RNA mixtures at highly imbalanced concentrations form tadpole-like structures that undergo a “tadpole–micelle–vesicle” transition, resulting in protein–RNA vesicles that resemble lipid membranes-like structures ^4^. Another example is provided by Knowles and colleagues ^5^, who demonstrated that rapid cooling of RNA–PEG co-condensates from elevated incubation temperatures induces the formation of unstable, biomolecule-depleted droplets within the condensates, driven by dynamically arrested phase separation.

While these studies have explored specific physical mechanisms underlying the formation of hollow condensates, a general thermodynamic driving force behind the formation of hollow condensates, if any, remains an open question. Here, we focus on a distinct phase separation phenomenon observed in double-stranded DNA (dsDNA)– protein interactive co-condensates (DPICs), wherein neither the protein nor dsDNA alone undergoes phase separation. We investigated the human transcription factor p53, specifically a truncated variant lacking the amino-terminal transactivation domain (TAD) termed p53^4M^ ΔTAD, which forms DPICs when combined with dsDNA ^6^. The p53^4M^ mutant contains four stabilizing mutations in the DNA-binding domain (M133L/V203A/N239Y/N268D), which confer enhanced thermodynamic stability and are commonly used to facilitate *in vitro* purification by preventing degradation ^7^ (Supplementary Fig. 1a). When mixed with 400-base pair (bp) dsDNA containing non-specific sequences (hereafter referred to as Random DNA), p53^4M^ ΔTAD forms droplet-like DPICs (Supplementary Fig. 1c(i), (ii), and (iv)). By contrast, incubation with 400-bp dsDNA bearing three p21-binding motifs (p21 DNA) (**Supplementary Information**) results in the formation of “pearl chain”-like condensates (Supplementary Fig. 1c(iii) and (v)). The p21 DNA motif is a well-characterized, high-affinity p53-binding site that plays a critical role in p53-mediated transcriptional activation and cell cycle arrest ^8,9^. Consistently, we found that p53 exhibits a stronger binding affinity for p21 DNA compared to Random DNA in *in vitro* conditions (Supplementary Fig. 1b).

In this study, we show that the addition of p21 DNA to droplet-like DPICs formed by p53^4M^ ΔTAD and Random DNA triggers a morphological transition from a biomolecule-rich condensate to a hollow architecture. *In vitro* droplet assays indicate that this transition is accompanied by p21 DNA–induced localized gelation at the condensate periphery. Guided by these experimental findings, we developed a three-component phase-field model that quantitatively captures the emergence of hollow condensates. Our simulations reveal that peripheral gelation driven by p21 DNA leads to a gradual depletion of protein and Random DNA in the condensate core. This redistribution drives spinodal decomposition, ultimately giving rise to a hollow structure.

## Results

### p53^4M^ ΔTAD can form three-dimensional hollow condensates with two types of dsDNA substrates

Following the formation of droplet-like DPICs by p53^4M^ ΔTAD and Random DNA (Fig. 1a and b(i)), the subsequent addition of p21 DNA induced a morphological transition to hollow condensates after 120 minutes of incubation (Fig. 1a and b(ii)). Notably, p53^4M^ ΔTAD and p21 DNA were also capable of forming “pearl chain”-like condensates (Supplementary Fig. 1c(iii) and (v)) ^6^. Confocal microscopy revealed that these hollow condensates are three-dimensional in nature (Fig. 1c). As a control, when p21 DNA was substituted with Random DNA, the condensates retained their biomolecule-rich morphology (Supplementary Fig. 2). Fluorescence intensity analysis further showed that the inner lumen and the surrounding dilute phase of the hollow condensates exhibited comparable fluorescence signals for both protein and Random DNA, markedly lower than the dense phase observed in biomolecule-rich DPICs (Supplementary Fig. 3a). Moreover, fluorescence correlation spectroscopy (FCS) measurements demonstrated that the diffusion coefficients of Random DNA in the lumen and the dilute phase were nearly identical (Supplementary Fig. 3b). Together, these findings indicate that the lumen and the surrounding dilute phase share similar molecular composition and mobility, confirming that the observed structures are indeed hollow condensates.

**Figure 1.**
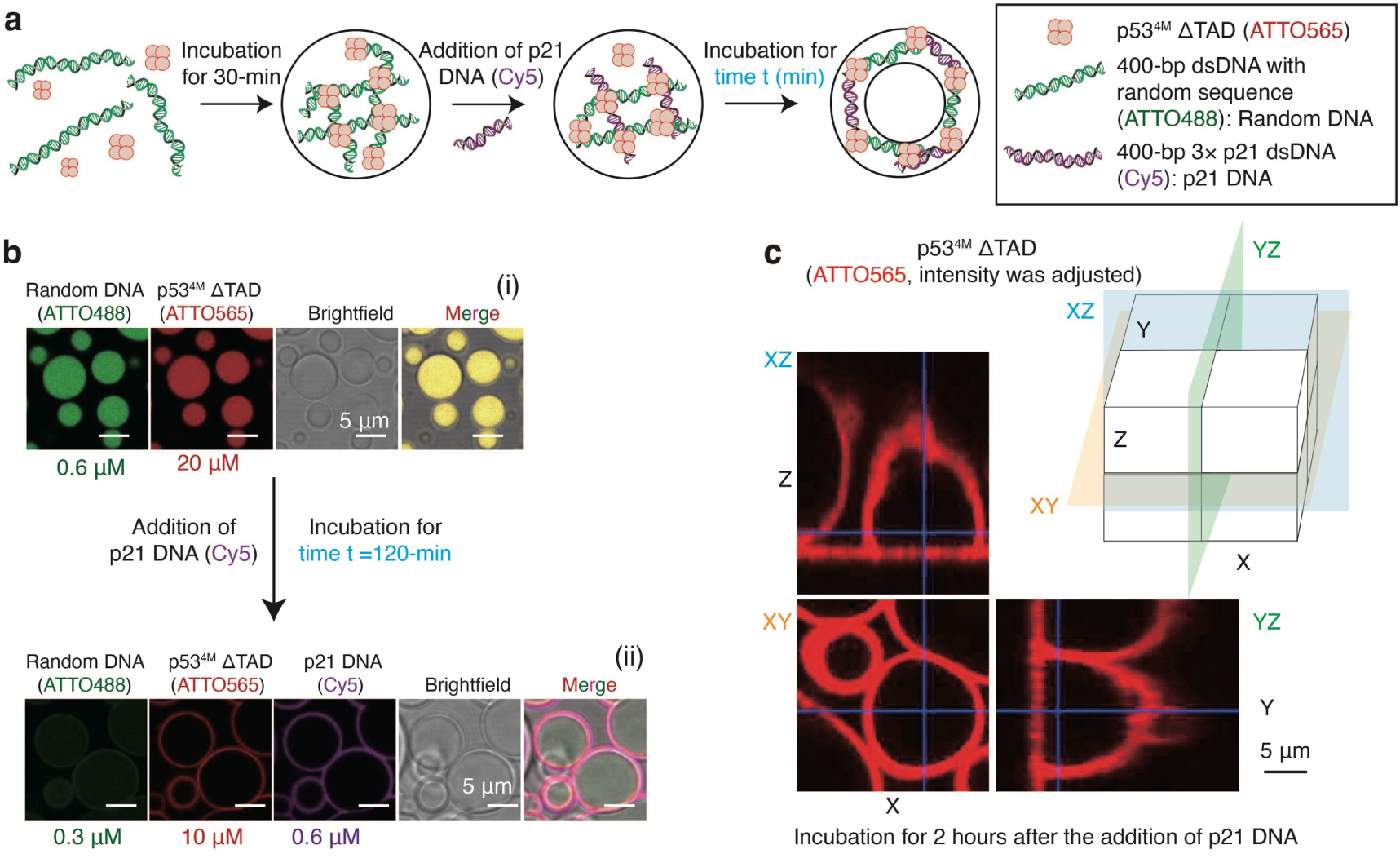
p53^4M^ ΔTAD forms three-dimensional hollow condensates with two types of double-stranded DNA substrates. (**a**) Schematic of the experimental design. (**b**) Experimental demonstration of hollow DPIC formation: (i) In the initial step, 20 μM ATTO565-labeled p53^4M^ ΔTAD and 0.6 μM ATTO488-labeled 400-bp Random DNA were mixed and incubated for 30 minutes at room temperature to form dual-color DPICs; (ii) After a 120-minute incubation following the addition of Cy5-labeled 400-bp p21 DNA, the final concentrations of p53^4M^ ΔTAD, Random DNA, and p21 DNA were adjusted to 10 μM, 0.3 μM, and 0.6 μM, respectively. (**c**) Three-dimensional reconstruction of hollow condensates shown in b(ii), displaying only the fluorescence signal from ATTO565-labeled p53^4M^ ΔTAD. *In vitro* droplet experiments were independently repeated three times (n = 3). The assay buffer consisted of 8 mM Tris-HCl (pH 7.5), 120 mM NaCl, 4% glycerol, and 16 mM DTT, without the use of crowding agents.

### The formation of hollow condensates is characterized by an abrupt dynamic transition

To investigate the dynamics of hollow condensate formation, we monitored structural transitions over time at t = 2, 20, 26, 27, 28, 29, 40, and 120 minutes, which corresponded to the incubation time after p21 DNA addition (Fig. 2a(i)-(ii) and Supplementary Movie 1). The corresponding fluorescence intensity profiles across the condensates are shown in Fig. 2a(iii). Interestingly, when hollow condensates are formed, p21 DNA is consistently enriched at the outer surface of the hollow condensate shell, whereas Random DNA localizes to the inner surface. Quantitative analysis of the fluorescence intensities of Random DNA and protein at the center of the DPICs revealed three distinct dynamic stages (Fig. 2b). Stage I was characterized by the diffusion and accumulation of p21 DNA at the periphery of biomolecule-rich, droplet-like DPICs, accompanied by a gradual depletion of protein and Random DNA in the condensate core. We defined a characteristic time parameter, τ_1_, to describe this initial phase (**Methods**). Stage II corresponded to an abrupt dynamic transition, during which protein and Random DNA intensities at the condensate center dropped sharply within a ∼2-minute interval. This phase was characterized by a second time constant, τ_2_ (**Methods**). Stage III reflected the stabilization of biomolecule-depleted state at the center of DPICs. Throughout the entire dynamical process, we also observed a marked increase in condensate size (Supplementary Fig. 4) and a process of protein and Random DNA escaping into the surrounding dilute phase (Supplementary Fig. 5), coinciding with the emergence of the hollow architecture.

**Figure 2.**
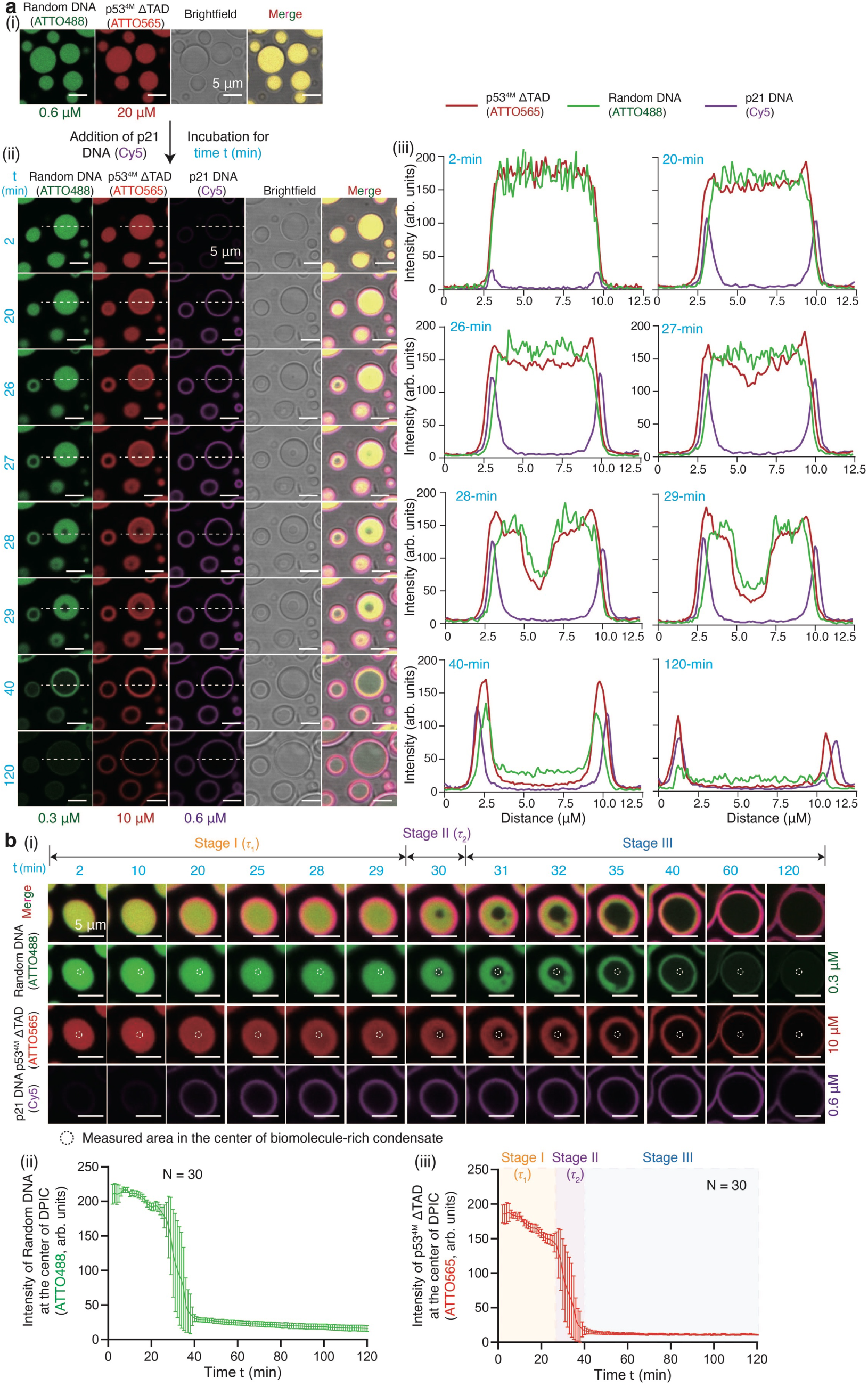
Formation of hollow condensates proceeds through three distinct dynamic stages. (**a**) Time-resolved visualization of hollow condensate formation. (i) Initially, 20 μM ATTO565-labeled p53^4M^ ΔTAD and 0.6 μM ATTO488-labeled 400-bp Random DNA were mixed and incubated for 30 minutes at room temperature, yielding dual-color, droplet-like DPICs; (ii) Cy5-labeled 400-bp p21 DNA was then added, and incubation continued at room temperature. Final concentrations were adjusted to 10 μM p53^4M^ ΔTAD, 0.3 μM Random DNA, and 0.6 μM p21 DNA. t denotes the elapsed time following p21 DNA addition. Representative fluorescence images at t = 2, 20, 26, 27, 28, 29, 40, and 120 minutes are shown; (iii) Fluorescence intensity profiles along the dashed line in panel a(ii) are plotted for each time point, showing the spatial distribution of p53^4M^ ΔTAD (red), Random DNA (green), and p21 DNA (purple). Independent in vitro droplet experiments were repeated three times (n = 3). (**b**) (i–iii) Temporal changes in fluorescence intensities of Random DNA and p53^4M^ ΔTAD in the central region of biomolecule-rich condensates. Data were collected from 30 individual condensates within a single experiment (N = 30). Error bars indicate mean ± s.d.

### The key factors to regulate the hollow condensate formation

We next asked what factors govern the formation of hollow condensate. We first increased the salt concentration from 120 mM to 500 mM, resulting in the complete dissolution of all hollow condensates (Supplementary Fig. 6). This finding suggests that their assembly is primarily driven by electrostatic interactions between DNA and protein, consistent with the mechanism underlying biomolecule-rich condensates ^6^.

We then investigated the impact of the sequential order of component addition. Hollow condensates formed only when protein and Random DNA were combined first, followed by the addition of p21 DNA. Conversely, no hollow structures were observed when protein was initially mixed with p21 DNA before adding Random DNA (Supplementary Fig. 7a), or when all three components were combined simultaneously (Supplementary Fig. 7b). These results highlight the critical role of assembly pathways in determining the final condensate architecture, suggesting the formation of hollow condensates has a kinetic origin.

We also examined the influence of p21 DNA concentration on the formation of hollow condensates. When the final p21 DNA concentration was below 0.225 μM (Fig. 3a(i)-(iii)), no hollow condensates were observed even after 120 minutes of incubation. In contrast, hollow condensates consistently formed at concentrations exceeding this threshold (Fig. 3a(iv)-(vii)). Thus, these findings indicate that a critical concentration of p21 DNA is necessary to drive the formation of hollow condensates. One should note that the threshold concentration of 0.225 μM p21 DNA is only applicable for a fixed incubation time of 120 minutes; the threshold may be lower if the incubation time is extended. For each experimental condition with different p21 DNA concentrations, we can measure the characteristic time parameters, τ_1_ and τ_2_, representing Stage I and II in Fig. 2b. We observed that τ_1_ displays a strong dependence on p21 DNA concentration (Fig. 3b(i)), whereas τ_2_ remains relatively stable across the tested concentration range (Fig. 3b(ii)).

**Figure 3.**
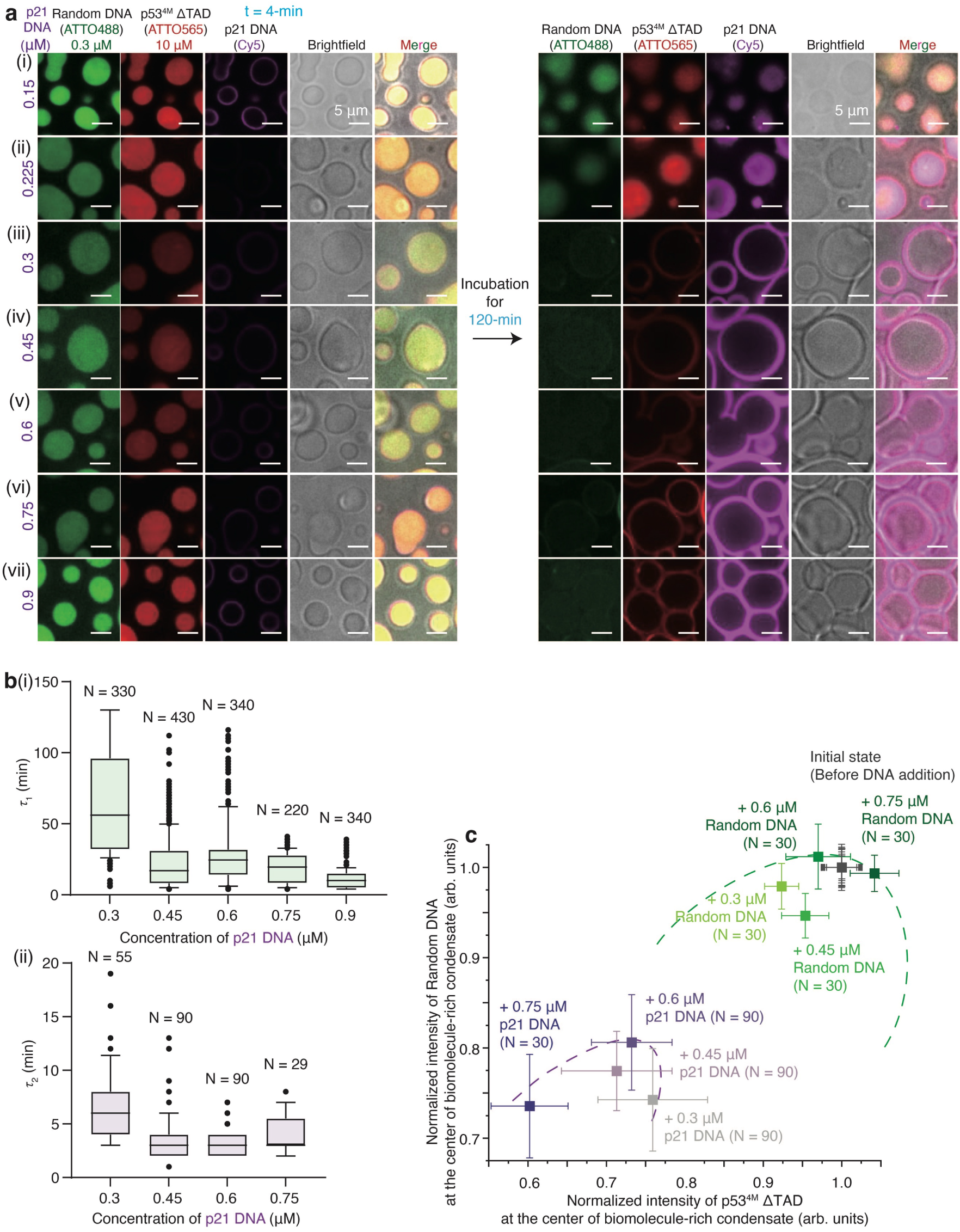
Concentration-dependent effects of p21 DNA on hollow condensate formation. (**a**) Droplet-like DPICs were initially formed by mixing 20 μM ATTO565-labeled p53^4M^ ΔTAD with 0.6 μM ATTO488-labeled Random DNA and incubating for 30 minutes at room temperature. Subsequently, Cy5-labeled p21 DNA was added at varying concentrations and incubated for an additional 120 minutes: (i) 0.15 μM; (ii) 0.225 μM; (iii) 0.3 μM; (iv) 0.45 μM; (v) 0.6 μM; (vi) 0.75 μM; (vii) 0.9 μM. Representative fluorescence images at incubation time t = 4-min and t = 120-min are shown. Independent *in vitro* droplet experiments were repeated three times (n = 3). (**b**) Boxplot of characteristic time constants τ_1_ and τ_2_ for p21 DNA concentrations ranging from 0.3 to 0.9 μM. N indicates the number of individual biomolecule-rich condensates analyzed under each condition. In box plots, the black line denotes the median, box edges represent the 25^th^ and 75^th^ percentiles, whiskers indicate the range excluding outliers, and outliers are shown as individual dots (•). (**c**) Phase diagram showing normalized fluorescence intensities of ATTO565-labeled p53^4M^ ΔTAD and ATTO488-labeled Random DNA at the center of condensates under increasing concentrations of p21 DNA (0.3, 0.45, 0.6, and 0.75 μM). Values are shown both before p21 DNA addition and at the end of Stage I. Control experiments in which Random DNA was used in place of p21 DNA are also included. Error bars indicate mean ± s.d. Green dashed lines mark the estimated binodal boundary, and purple dashed lines represent the spinodal boundary, as confirmed by our phase-field model (see Figs. 5–6).

Interestingly, under all experimental conditions with varying concentrations of p21 DNA (0.3, 0.45, 0.6, and 0.75 μM), the fluorescence intensities of Random DNA and protein at the center of biomolecule-rich condensates decreased during Stage I (Fig. 3c), which means that the concentrations of Random DNA and protein shifted from the concentrations of the dense phase toward lower concentrations, triggering spinodal decomposition as we show later. As a control, replacing p21 DNA with Random DNA under identical conditions resulted in negligible shifts.

### The hollow condensate formation accompanies the process of gelation

Building on our previous findings that droplet-like DPICs composed of protein and Random DNA exhibit distinct viscoelastic properties ^6^, we next examined the material properties of the hollow condensates. Fluorescence recovery after photobleaching (FRAP) experiments revealed that the shell regions of hollow condensates exhibited markedly lower and slower recovery of p53^4M^ ΔTAD compared to that inside biomolecule-rich condensates (Fig. 4a). To further assess the mechanical properties, we performed atomic force microscopy-based force spectroscopy (AFM-FS), which demonstrated that DPICs treated with p21 DNA displayed significantly higher Young’s modulus values than those incubated with Random DNA (Fig. 4b). Together, the FRAP and AFM-FS experiments suggest that the addition of p21 DNA induces a transition from a liquid-like to a gel-like material state during the hollow condensate formation.

**Figure 4.**
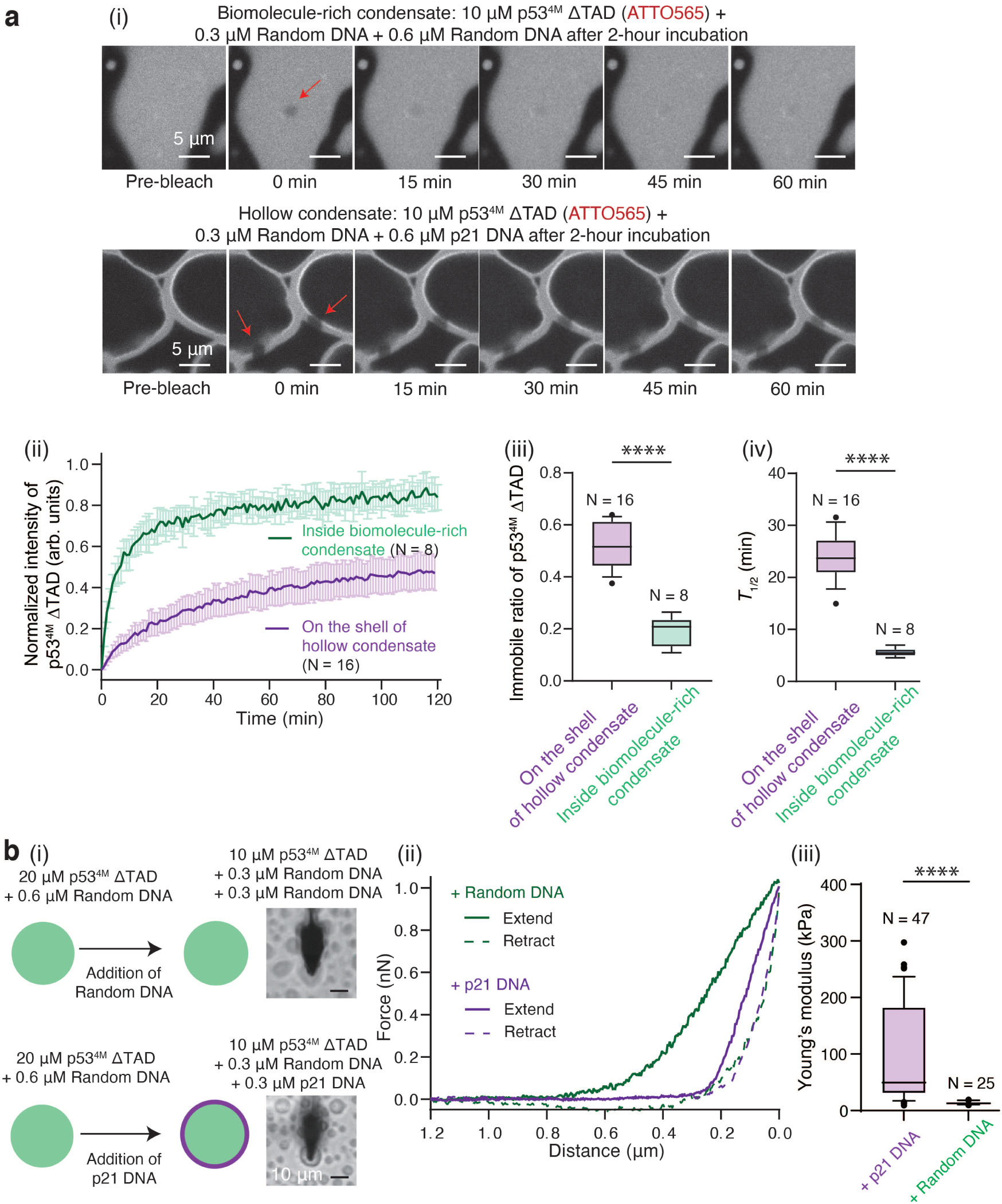
Hollow condensate formation is accompanied by gelation. (**a**) FRAP analysis of condensate dynamics. (i) Droplet-like DPICs were generated by incubating 20 μM ATTO565-labeled p53^4M^ ΔTAD with 0.6 μM Random DNA for 30 min at room temperature. Subsequently, either additional Random DNA (Condition 1) or p21 DNA (Condition 2) was added, yielding final concentrations of 10 μM p53^4M^ ΔTAD, 0.3 μM Random DNA, and 0.6 μM of the added DNA. Samples were incubated for 2 hours prior to photobleaching, followed by a 2-hour recovery period. Red arrows indicate bleached regions. (ii) FRAP recovery curves for Conditions 1 and 2. Data represent mean ± s.d. (N = 16 for Condition 1; N = 8 for Condition 2). (iii–iv) Box plots showing mobile fraction (iii) and recovery half-time (*T*_1/2_) (iv) for both conditions. (**b**) AFM-FS measurements of condensate mechanics. (i) Schematic of AFM-FS assay. Initial DPICs were prepared by incubating 20 μM p53^4M^ ΔTAD with 0.6 μM Random DNA for 20 minutes. Random DNA (Condition 3) or p21 DNA (Condition 4) was then added and incubated for an additional 20 minutes. Final concentrations were 10 μM p53^4M^ ΔTAD, 0.3 μM Random DNA, and 0.3 μM added DNA. (ii) Representative force– distance curves for Conditions 3 (green) and 4 (purple). Solid lines represent extension phases; dashed lines represent retraction. (iii) Box plot of Young’s modulus for Conditions 3 (N = 25) and 4 (N = 47). Independent *in vitro* droplet experiments in (a) and (b) were repeated twice. In box plots, the center line indicates the median; box edges denote the 25^th^ and 75^th^ percentiles; whiskers represent the range excluding outliers, shown as dots (•). Statistical comparisons were performed using unpaired two-tailed t-tests. P value style: GP: 0.1234 (ns), 0.0332 (*), 0.0021 (**), 0.0002 (***), < 0.0001 (****). Error bars represent mean ± s.d.

### A continuum model of phase separation recapitulates the hollow condensate formation

To elucidate the physical mechanism underlying the formation of hollow condensates, we sought to develop a continuum model. Our experimental system comprises three biomolecular components: protein (p53^4M^ ΔTAD), Random DNA, and p21 DNA. Notably, none of these components undergoes phase separation in isolation, and the only attractive interactions present are between protein and Random DNA, and between protein and p21 DNA. Building on these experimental observations, we formulated a three-component continuum model (**Supplementary Methods**). We assumed that the system follows Cahn-Hilliard dynamics, governed by a free energy driving the condensate formation between p53^4M^ ΔTAD and Random DNA or p21 DNA ^10–14^ (Fig. 5a).

**Figure 5.**
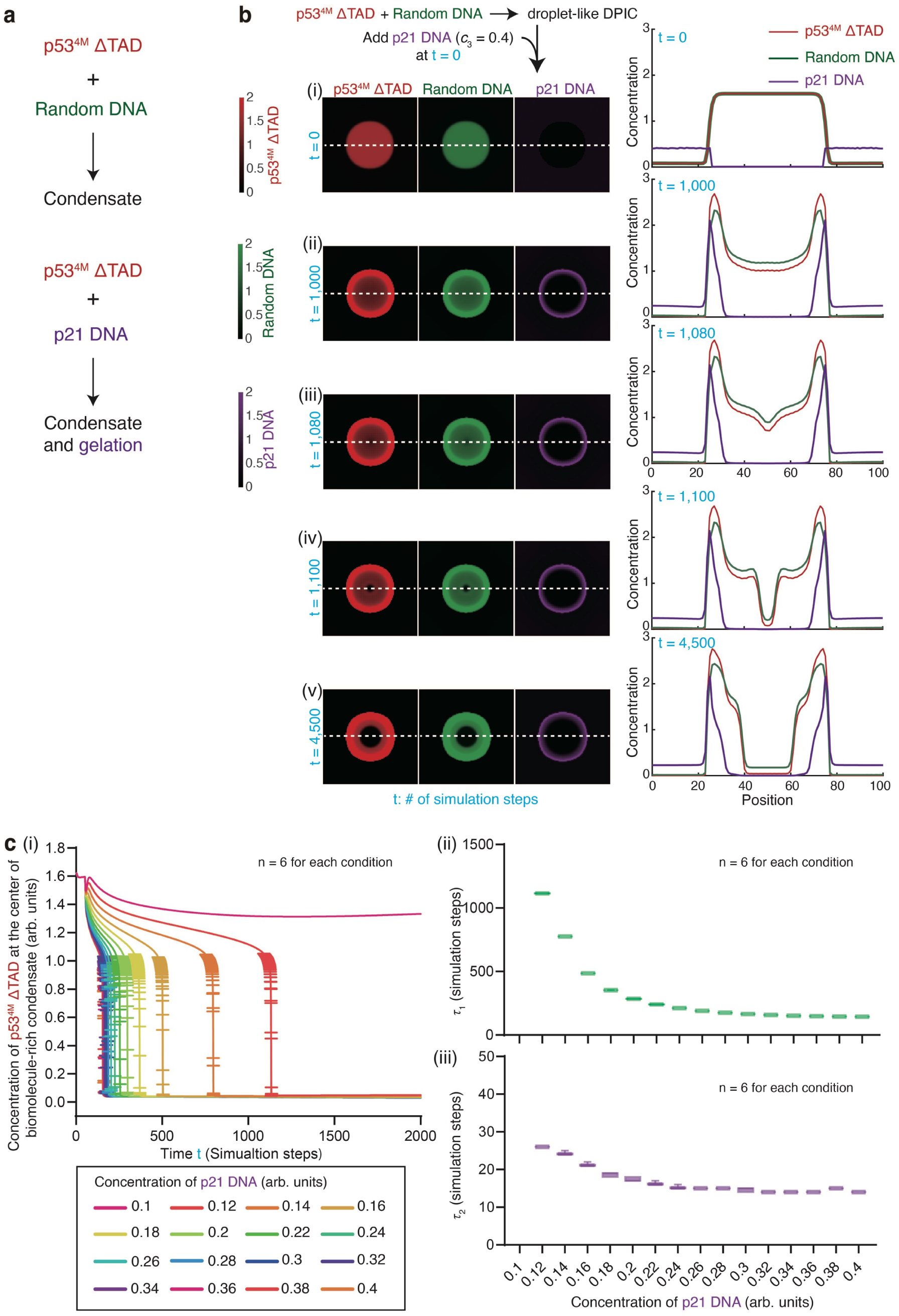
A continuum phase-field model recapitulates hollow condensate formation. (**a**) Schematic illustrating the initial conditions of the three-component continuum phase-field model. (**b**) Simulated time course of hollow condensate formation. Heatmaps show the spatial distribution of protein, Random DNA, and p21 DNA at selected time points, alongside intensity profiles along the white dashed line. In this simulation, p21 DNA was introduced at a concentration of *c*_3_ = 0.4, and the initial condensate radius was set to *R* = 25 (see **Supplementary Methods**). (**c**) The model captures three distinct dynamic stages of hollow condensate formation. (i) Temporal evolution of protein (p53) concentration upon addition of varying concentrations of p21 DNA. (ii–iii) Quantification of the characteristic time constants τ_1_ (ii) and τ_2_ (iii). Each simulation in panel (c) was independently repeated six times (n = 6). Error bars represent mean ± s.d.

Fluorescence signals in our *in vitro* droplet assays revealed that p21 DNA is consistently enriched at the outer surface of the hollow condensate shell (Fig. 2a). Notably, adjacent hollow condensates fail to undergo fusion (Supplementary Movie 1), and both FRAP and AFM-FS measurements also indicate that hollow condensate formation is accompanied by a transition to a gel-like state (Fig. 4). These observations suggest that p21 DNA induces localized gelation at the condensate periphery, which in turn drives the formation of hollow structures. To capture this behavior in our model, we incorporated a gel field that emerges when the local concentrations of both protein and p21 DNA exceed a critical threshold (Fig. 5a and **Supplementary Methods**). This gelation field significantly impedes molecular diffusion, consistent with theoretical predictions ^15^.

To initiate the numerical simulation, we first generated a protein–Random DNA condensate as the initial condition. Following the experimental procedure (Fig. 2a), we initiated the simulation by introducing a nearly uniform concentration of p21 DNA in the external dilute phase, defining this point as time zero (t = 0 step; Fig. 5b(i)). In the model, the concentration of p21 DNA was denoted as *c*_3_ (see **Supplementary Methods**), with an initial value of *c*_3_ = 0.4. The full simulation, spanning t = 4,500 steps, is presented in Supplementary Movie 2. The representative snapshots of t = 1,000, 1,080, 1,100, and 4,500 and the corresponding component concentration profiles across the condensates are shown in Fig. 5b(ii)-(v). Consistent with experimental observations, the simulation revealed three distinct stages of hollow condensate formation. Stage I (t < 1,000 steps) was characterized by the diffusion and accumulation of p21 DNA around the protein–Random DNA condensates, and a gradual decline in the concentrations of protein and Random DNA in the central region of the condensate (Fig. 5b(ii)); Stage II (t = 1,000–1,100 steps) marked an abrupt dynamic transition (Fig. 5b(iii)-(iv)), during which protein and Random DNA intensities at the condensate center dropped sharply within a ∼100-steps interval); Stage III (t > 1,100 steps) reflected the stabilization of the hollow lumen, which maintained dilute-phase properties over extended periods.

To validate our model, we first repeated the simulation in Fig. 5b, replacing p21 DNA with Random DNA. Even after 4,500 simulation steps, hollow condensates failed to form (Supplementary Fig. 8a and Supplementary Movie 3), consistent with our experimental observations (Supplementary Fig. 2). Given that reducing p21 DNA concentration inhibited hollow condensate formation in our experiments (Fig. 3a), we next repeated the simulation in Fig. 5a and decreased the p21 DNA concentration (*c*_3_) from 0.4 to 0.1 (**Supplementary Methods**). In agreement with experiments, no hollow condensates formed after 4,500 simulation steps (Supplementary Fig. 8b and Supplementary Movie 4). To further examine the effect of p21 DNA concentration, we systematically varied *c*_3_from 0.1 to 0.4 in increments of 0.02. The same component intensity analyses (Fig. 2b) yielded the simulated characteristic time parameters, τ_1_ and τ_2_ (Fig. 5c), which closely matched experimental results (Fig. 3b). These findings demonstrate that our continuum model successfully reproduced the key experimental observations (Figs. 1-3).

We next investigated whether our model could elucidate the physical mechanism underlying the formation of hollow condensates. We performed numerical simulations to track the temporal evolution of component concentrations and thermodynamic stability within the central region of DPICs during the emergence of hollow structures (**Supplementary Methods**). In this analysis (Fig. 6a), we computed the Hessian matrix of the free energy, defined as (*H*)_*ij*_ = ∂^2^*f*/ ∂*c*_*i*_ ∂*c*_*i*_, using the simulated concentrations of p53 and Random DNA obtained from the model presented in Fig. 5. The contribution of p21 DNA was excluded, given its negligible concentration at the center of DPIC. Temporal profiles of p53 and Random DNA concentrations are shown as red and green lines, respectively (Fig. 6a). From these simulation results, we calculated the free energy from the core region of the condensate, and derived the minimum eigenvalue of the Hessian matrix (Min Eigenvalue), shown as the blue line. This eigenvalue serves as a proxy for local thermodynamic stability in the condensate core: values significantly greater than zero indicate a stable state, whereas values below zero reflect an unstable state.

**Figure 6.**
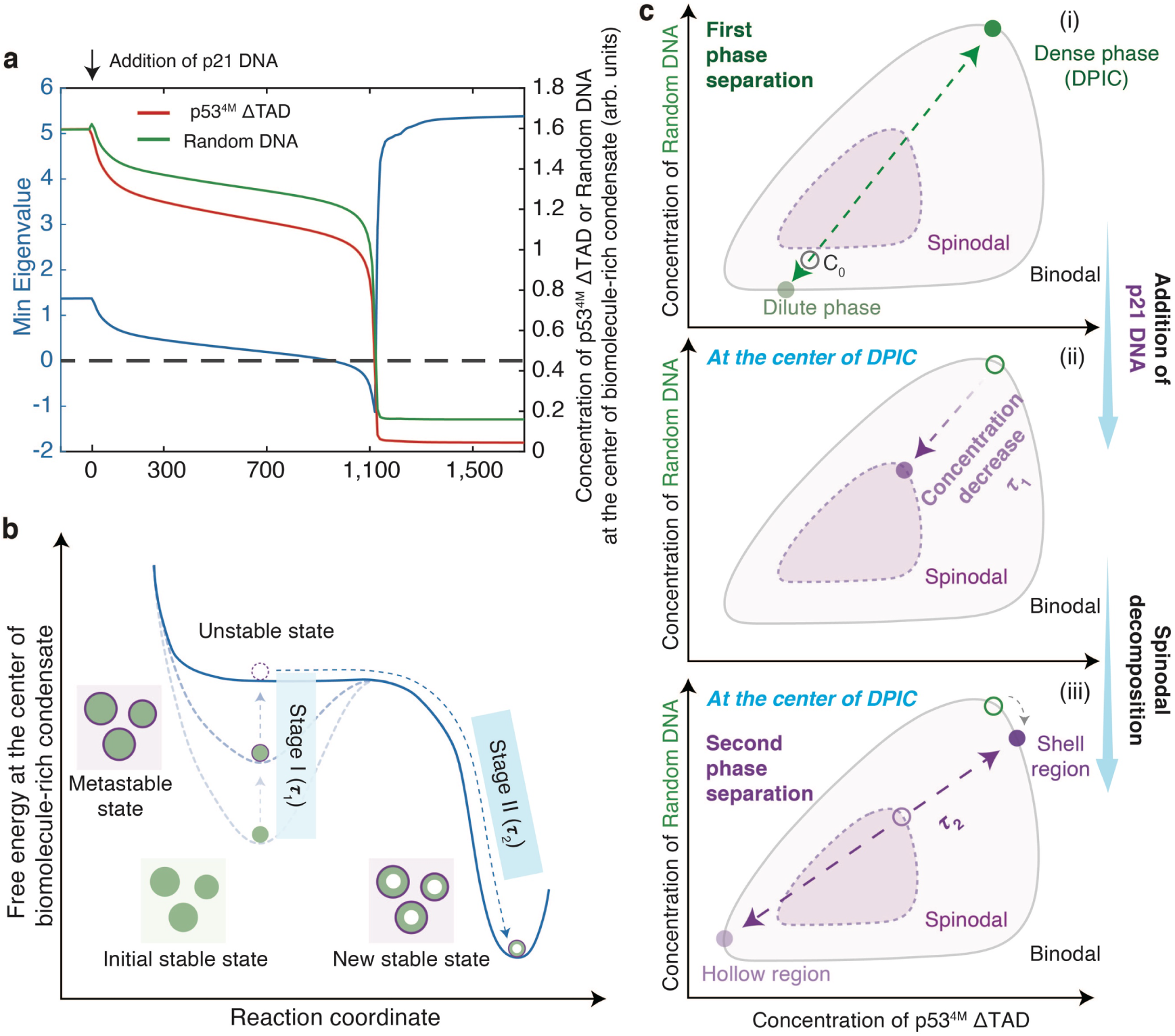
Mechanistic insights from modeling reveal the physical basis of hollow condensate formation. (**a**) Temporal evolution of component concentrations and thermodynamic stability at the center of biomolecule-rich condensates in numerical simulations. The red and green curves represent the concentrations of protein and Random DNA, respectively. The blue curve denotes the Min Eigenvalue of the Hessian matrix, reflecting local thermodynamic stability. (**b**) Schematic representation of the energy landscape underlying hollow condensate formation. During Stage I (τ_1_), p21 DNA perturbs the system, driving the condensate from a metastable to an unstable state. In Stage II (τ_2_), spinodal decomposition ensues, leading to a new stable state characterized by a condensed shell and an internal dilute phase. (**c**) Proposed physical mechanism of hollow condensate formation. (i) p53^4M^ ΔTAD initially co-phase separates with Random DNA, producing a dense biomolecule-rich condensate (DPIC) surrounded by a dilute phase. (ii–iii) Upon addition of p21 DNA, the system undergoes a secondary transition. (ii) Protein and Random DNA concentrations in the condensate core progressively decline; (iii) Once local concentrations cross the spinodal threshold, internal phase separation is triggered, yielding a hollow architecture with a condensed shell and a central biomolecule-depleted region. The hollow circle denotes the initial concentration, while the filled circle indicates the final concentration.

These analyses captured the dynamic evolution of the central region of the DPIC (Fig. 6a). Before the addition of p21 DNA, the concentrations of both p53 protein and Random DNA remained high, and the Min Eigenvalue was well above zero—indicating a stable biomolecule-rich phase. Upon the introduction of p21 DNA (t = 0–1,000 steps), both concentrations in the central region gradually declined as they flowed toward the condensate periphery due to the attractive interaction between the protein and p21 DNA (Fig. 5b), consistent with experimental observations during Stage I (Fig. 2b). During this period, the Min Eigenvalue also decreased but remained positive, suggesting that the system had transitioned into a metastable state. Between t = 1000 and 1100 steps, a sharp drop in both molecular concentrations coincided with the Min Eigenvalue crossing below zero, which means the onset of spinodal decomposition within the condensate core, indicating a second phase separation. This transition aligns with experimental observations corresponding to Stage II (Fig. 2b). Finally, after t > 1,100 steps, both concentrations stabilized at low levels, and the Min Eigenvalue returned to a high positive value, indicating that the central region had entered a new, thermodynamically stable state characterized by biomolecular depletion—consistent with Stage III observed experimentally (Fig. 2b).

Together, these experimental findings and our model suggest that p21 DNA induces localized gelation at the condensate periphery, leading to a progressive depletion of protein and Random DNA in the core. This compositional shift drives spinodal decomposition inside the condensate, ultimately giving rise to hollow condensates.

## Discussion

In this study, we demonstrate that the addition of p21 DNA to droplet-like DPICs triggers a morphological transition from biomolecule-rich condensates to a hollow architecture (Fig. 1b). This transformation proceeds through three kinetically distinct stages (Fig. 2b). Stage I, characterized by a timescale τ_1_, corresponds to the diffusion and accumulation of p21 DNA around the biomolecule-rich condensate. Stage II, defined by the characteristic time τ_2_, marks a rapid and abrupt transition. Stage III represents the stabilization of a biomolecule-depleted lumen, which persists over extended time scales. We further identified salt concentration (Supplementary Fig. 6), the order of component addition (Supplementary Fig. 7), and the concentration of p21 DNA (Fig. 3a-b) as key parameters governing hollow condensate formation. Notably, τ_1_ exhibited strong dependence on p21 DNA concentration, while τ_2_ remained largely unaffected across conditions. The *in vitro* droplet assay, FRAP and AFM-FS analyses revealed that the formation of hollow condensates is accompanied by a gelation process at the condensate periphery (Fig. 4). Guided by these findings, we constructed a three-component continuum model incorporating the only hypothesis of gelation field to represent localized p21 DNA-induced gelation (Fig. 5). This framework quantitatively recapitulates the key experimental observations presented in Figs. 1–3. Using this model, we analyzed the Min Eigenvalue of the free energy Hessian matrix as a proxy for local stability within the condensate core (Fig. 6a).

These analyses allowed us to pinpoint the onset of internal phase instability, offering mechanistic insight into the emergence of hollow morphology, which is schematically represented by an energy landscape (Fig. 6b). Prior to the addition of p21 DNA, protein and Random DNA assemble into stable, biomolecule-rich DPICs. Upon p21 DNA introduction, a gradual decline in the concentrations of these components at the center of DPICs drives the system into a metastable state, and then pushes the DPICs to reach an unstable state—corresponding to Stage I (τ_1_). The instability then triggers abrupt decreases of the concentrations of protein and Random DNA at the center of DPICs, though spinodal decomposition—corresponding to Stage II (τ_2_). This process ultimately leads to the formation of a new, thermodynamically favorable state characterized by a core depleted of biomolecules (Stage III).

By integrating the proposed energy landscape with our experimental observations (Fig. 3c), we further delineate a physical mechanism underlying the formation of hollow structures. A schematic two-component phase diagram is presented in Fig. 6c to illustrate this process. Initially, a primary phase separation partitions proteins and Random DNA into a dense phase (DPICs) and a surrounding dilute phase (Fig. 6c(i)). Upon the addition of p21 DNA, during Stage I, the internal concentrations of protein and Random DNA within the DPIC decrease, shifting from the dense-phase concentrations at the binodal boundary to lower values (Fig. 6c(ii)). This transition facilitates a second phase separation within the condensate, producing a new internal dilute phase corresponding to the hollow core (Fig. 6c(iii)). Interestingly, this transition involves spontaneous destabilization of the condensate interior, strongly suggesting that the component concentrations reach the spinodal region of the phase diagram during Stage I—a hypothesis supported by our theoretical calculations (Fig. 6a) and the quantitative agreement between our simulations and experiments (Fig. 3c). This behavior is reminiscent of spinodal decomposition phenomena reported in other biomolecular condensate systems ^5,14,16,17^.

Our findings highlight several compelling directions for future investigation. First, the current model does not capture the pronounced increase in condensate size observed during the transition from a homogeneous to a hollow architecture (Supplementary Fig. 4), likely due to its omission of mechanical details. In particular, the simplified gel field does not incorporate the constitutive behavior of the gelled interface. A more advanced framework—potentially one that integrates fluid–structure interactions—may be necessary to accurately describe this growth process. In parallel, the current model does not recapitulate the experimentally observed escape of protein and Random DNA into the surrounding dilute phase (Supplementary Fig. 5), likely due to its omission of microscale features of the gelled interface—such as dynamic changes in pore size within the elastic network—that may facilitate this release. An agent-based modeling framework may provide an alternative approach to capture and further elucidate these compositional dynamics.

Second, our findings highlight the critical role of assembly pathways in shaping the ultimate architecture of biomolecular condensates (Supplementary Fig. 7). In living cells, condensates are typically composed of multiple dynamic components whose identities and concentrations evolve over time. The composition of emerging condensates in living cells is highly sensitive to the local molecular environment. Variations in the order of component addition can result in condensates of distinct compositions that become trapped in different metastable states, potentially conferring unique biological functions. Thus, biomolecular condensates may never reach the thermodynamic equilibrium state under physiological conditions. Instead, they may remain kinetically arrested in non-equilibrium states that reflect their assembly history. This principle has important implications for understanding the functional diversity of condensates and merits further investigation in future studies.

Third, prior studies have highlighted the functional potential of hollow condensates in molecular encapsulation and delivery. For example, protamine–RNA hollow condensates have been demonstrated to selectively enrich single-stranded DNA (ssDNA) and dsDNA within their lumens ^4^. More recently, hollow condensates formed from cholesterol-modified DNA and histones can be harnessed for drug delivery, encapsulating biologics such as viral particles and mRNA conjugated to nucleic acids or polypeptides ^18^. These findings collectively suggest that dsDNA-p53-based hollow condensates may similarly serve as versatile carriers for nucleic acid-linked therapeutics. Exploring this possibility represents a compelling direction for future research.

In summary, our findings elucidate the key biophysical mechanisms underlying the formation of three-component hollow condensates. These insights lay the groundwork for the rational modulation of biomolecular condensation, establishing a foundation for leveraging condensate morphologies in therapeutic intervention in human diseases and biotechnologies.

## Author Contributions

C.L. prepared biological samples, conducted all experiments, performed data analysis, and wrote the manuscript. L.M. and J.L. built up the theoretical model and wrote the manuscript. Y.T. assisted C.L. Z.Q. supervised the project, experimental designs, and data analysis, and wrote the manuscript with input from all authors.

## Acknowledgments

We thank the National Center for Protein Sciences at Peking University in Beijing, China, particularly Dr. Siying Qin, for technical help with AFM. We thank Dunjin Zheng, the engineer from LightEdge Technology Co., LTD for providing FCS testing. We thank Dr. Luhua Lai and Chun Tang (Peking University), Dr. Xiangze Zeng (Hong Kong Baptist University), and the members of the Z.Q. laboratory for comments on the manuscript.

## Data and code availability

The code of the mathematical model is available in https://github.com/LingyuMeng99/Hollow-Structure-Simulation.

## Methods

### Protein expression and purification

Based on our previous experimental results, the truncated p53 protein lacking the transcriptional activation domain (TAD) and carrying four mutations (M133L/V203A/N239Y/N268D), which was called p53^4M^ ΔTAD, cannot undergo phase separation alone. This truncated variant requires intermolecular interaction with dsDNA to form DPICs ^6^. Thus, we used p53^4M^ ΔTAD as a model system for *in vitro* droplet experiments in this study.

The expression plasmid of p53^4M^ ΔTAD was transformed into *E. coli* BL21(DE3) competent cells, followed by overnight incubation on LB agar plates at 37 °C. A single bacterial colony was selected and grown overnight in 5 mL of LB medium under shaking conditions (220 rpm) at 37 °C. Subsequently, the starter culture was transferred into 2 L of LB medium and allowed to grow until reaching an OD600 of 0.6. Protein expression was induced by the supplement of 0.3 mM IPTG and 0.1 mM ZnCl_2_, followed by 18-hour incubation at 16 °C with continuous shaking at 180 rpm.

The bacterial cells were harvested via centrifugation at 4,000× g and resuspended in pre-cold lysis buffer (25 mM Tris-HCl pH 7.5, 500 mM NaCl, 5 mM imidazole, 0.25‰ β-mercaptoethanol, 5% glycerol) supplemented with 1 mM PMSF. Cells were sonicated on ice, and then centrifugated at 18,000× rpm for 30 minutes. The supernatant was filtered and loaded to Ni-NTA resin, with sequential washing steps employing wash buffer (25 mM Tris-HCl (pH 7.5), 500 mM NaCl, 20 mM imidazole, 0.25‰ β-mercaptoethanol, 5% glycerol). The bounded protein was eluted by the elution buffer (25 mM Tris-HCl (pH 7.5), 500 mM NaCl, 300 mM imidazole, 0.25‰ β-mercaptoethanol, 5% glycerol). Final purification was achieved through gel filtration using a Superdex 200 Increase GL 10/300 column (GE Healthcare) equilibrated with storage buffer (20 mM Tris-HCl (pH 7.5), 300 mM NaCl, 10% glycerol, 40 mM DTT). Purified proteins were first analyzed by SDS-PAGE for verification, then flash-frozen in liquid nitrogen and stored at -80 °C for preservation.

### Fluorescence labeling of protein and DNA

To visualize the protein under confocal microscopy or microplate reader, p53^4M^ ΔTAD was conjugated with ATTO565 NHS ester (Sigma, Cat#72464) or AF647 NHS ester (DuoFluor, Cat#D10157) at a 1:2.5 molar ratio in 0.1 M NaHCO3 buffer through a 1-hour incubation at 4 °C. The labeled proteins underwent purification via gel filtration using a Superdex 200 Increase GL 10/300 column pre-equilibrated with storage buffer. The purified fluorescently tagged proteins were subsequently flash-frozen and archived at -80 °C for long-term storage.

The ATTO488-labeled 400-bp dsDNA containing non-specific sequence (Random DNA) and Cy5-labeled 400-bp dsDNA bearing three p21-binding motifs (p21 DNA) were obtained by PCR reaction. The template was a sequence containing no p53-specific binding sites on the genome of wild type Lambda phage. One primer carried an ATTO488 or Cy5 tag at the 5’ terminal, and the other primer did not carry any tag. The PCR product was purified by DNA Extraction Kit (Vazyme, #DC301), dissolved in ddH_2_O and then stored in -20 °C condition.

### Electrophoretic mobility shift assay (EMSA)

30-bp dsDNA probes were generated through thermal annealing of complementary primers. A 5’-Quasar670-labeled top strand and unlabeled bottom strand were combined at a 1:1.2 molar ratio in annealing buffer (40 mM Tris-HCl (pH 8.0), 50 mM NaCl, 10 mM MgCl₂), followed by denaturation at 95 °C for 5 minutes and gradual cooling to room temperature (RT) on the heater. Annealed probes were quantified as 1 μM stock solutions.

The EMSA experiments were performed by incubating serially diluted p53^4M^ ΔTAD with 0.05 μM DNA probes in p53 binding buffer (20 mM HEPES (pH 7.9), 150 mM NaCl, 2 mM MgCl_2_, 1 mM DTT, 0.5 mg/mL BSA). After 30-min incubation at RT, DNA-protein complexes were resolved on 1% agarose gels in TBE buffer under 60 V for 90 minutes at 4 °C. Gels were imaged using an Amersham Typhoon scanner (Cy5 channel) for EMSA analysis.

It should be emphasized that to reduce the impact of different purified batches of proteins on the experimental results, we used proteins with the EMSA activity which was similar to that in Supplementary Fig. 1 for experiments in this article.

### *In vitro* droplet experiment and data analysis

#### *In vitro* droplet assay

The stock solutions of protein and DNA were pre-cleared by centrifugation with 13,800× g for 10 minutes at 4°C. To achieve specific protein and DNA concentrations, proteins were diluted in storage buffer while DNA was diluted in ddH₂O. Protein-DNA mixtures were homogenized by repeated pipetting and transferred to a 384-well plate (Cellvis, #P384-1.5H-N). Following 30-min incubation at RT, phase-separated droplets were visualized using confocal microscopy (Leica TCS SP8, 100× objective). All droplet assays were performed in droplet buffer (8 mM Tris-HCl (pH 7.5), 120 mM NaCl, 4% Glycerol, and 16 mM DTT).

To investigate the formation of hollow condensates, the droplet buffer containing p21 DNA with different concentrations was introduced into the droplet systems. The mixture was gently pipetted for 5 times to homogenize the reaction system while minimizing the damage to the existed DPICs. Following incubation at RT for specified time intervals, samples were imaged by confocal microscopy.

The image data was analyzed by Fiji (ImageJ Version: 2.0.0-rc-61/1.52n). Fluorescence images were initially converted to 8-bit format. We drew a straight line across the region of interest (ROI) and used the Fiji (Analyze / Plot Profile) to extract the fluorescence intensity of each pixel along the line for the protein and DNAs.

Since fluorescence intensity is highly correlated with fluorescent molecule concentration, we used the average intensity of p53^4M^ ΔTAD and Random DNA inside DPICs to represent their concentrations. Using the Fiji (Analyze / Tools / ROI manager), we selected circular regions with a diameter of 1 μm as ROIs inside 30 independent DPICs in a single imaging experiment. The “multi-measure” tool was then used to record the time series in the average fluorescence intensity of proteins (ATTO565) and Random DNA (ATTO488) in all ROIs.

To characterize the area increasement of DPICs during hollow condensate formation, the image data of ATTO565 channel which represented ATTO565-labeled p53^4M^ ΔTAD was imported to Fiji. Auto threshold was applied to the frame displaying the weakest fluorescence signal via Fiji (Image / Adjust / Threshold), and this threshold setting was implemented to the entire image stack to convert them to a binary format. Next, Fiji (Analyze / Analyze Particles) was utilized with the “Include holes” option activated to ensure measurement of the whole area of DPIC at each time point.

#### Fluorescence recovery after photobleaching (FRAP)

To compare the dynamic properties of p53^4M^ ΔTAD on the periphery of hollow condensates or inside the biomolecule-rich condensates, two kinds of condensates were pre-incubated at RT for 2 hours to make the system as stable as possible. FRAP assays were conducted using a Leica SP8 confocal microscope equipped with a 100× objective. The size of the ROIs for bleaching was approximately 2 μm × 2 μm, and 5-10 different ROIs could be measured in one experiment. The power of 488 and 561-nm lasers were both set to 100%, and a total of 20 rounds of bleaching were performed. Then, the recovery was recorded at a frequency of 1 frame / min for 120 minutes.

The image data was imported to Fiji and adjusted to 8-bit format for analyze. To correct for photobleaching effects during the 120-min imaging period, the fluorescence intensity of each ROI was normalized to control region (*I*_*t*_*norm*_1_ = *I*_*t*_ / *I*_*t*_*control*_). Background correction was then performed by subtracting the fluorescence intensity of the first post-bleaching frame from all subsequent time points, effectively establishing a baseline value of zero (*I*_*t*_*norm*_2_ = *I*_*t*_*norm*_1_ − *I*_*t*1_*norm*_1_). Finally, the data were normalized to the pre-bleaching intensity to standardize the initial fluorescence level to 1 (*I*_*t*_*norm*_3_ = *I*_*t*_*norm*_2_ / *I_pre_norm_*__2_).

The normalized recovery curves of the ROIs were fitted using the equation *I*_*t*_*norm*_3_ = *A* · (1 − *e*^−τ*tt*^), where *A* is the mobile fraction of the protein, and τ is the recovery time constant. The immobile ratio of protein is 1 − *A* . The half-time of fluorescence recovery (*T*_1/2_) was then calculated using the equation 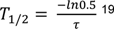.

#### Measurement of τ_1_ and τ_2_

For experiments measuring τ_1_, the preliminary workflow aligns with the previously described procedures for preparing biomolecule-rich condensate and introducing the p21 DNA-containing droplet buffer. After adding the p21 DNA, the sample was placed under a spinning-disk confocal microscope (UltraView VoX, 100× objective). Approximately 10 independent fields of view were selected and continuously imaged for 2 hours at a frame rate of 2 minutes per frame. The ATTO565 channel images, p53^4M^ ΔTAD, were imported into Fiji. Using Fiji (Image / Adjust / Threshold), an auto-threshold was applied to the frame with the weakest fluorescence signal, and this threshold was used to convert the whole image stack to a binary format. Subsequently, 10 independent DPICs were randomly selected in each observation field, and the frame numbers at which hollow structures appeared were manually recorded. The time corresponding to the preceding frame was defined as τ_1_.

For experiments measuring τ_2_, the sample preparing is the same as above. After adding the p21 DNA, the sample was placed under a Leica SP8 confocal microscope equipped with a 100× objective. Each experiment was continuously imaged for 2 hours at a frame rate of 1 minutes per frame. The fluorescence images of p53^4M^ ΔTAD were imported into Fiji, and approximately 30 independent DPICs were randomly selected in one field of view to record the changes in fluorescence intensity over time in their central regions. For cases where the initial average intensity was approximately 200 arb. units, the frame preceding the first frame where the change in the minimum intensity of the central region between two consecutive frames was ≥ 30 was defined as the onset of τ_2_, and the frame preceding the first frame where the change in the maximum intensity of the central region between two consecutive frames was ≤ 10 was defined as the offset of τ_2_. Based on this criterion, τ_2_ for each DPIC was manually determined.

For τ_1_ and τ_2_ in the numerical simulation, we observed that the protein concentration remains in a relatively stable state (Stage A) before decreasing to 1.0, and enters another stable state (Stage B) after decreasing to 0.05. Therefore, we define the duration of Stage A as τ_1_, and the time interval between Stage B and Stage A as τ_2_.

#### Measurement of the release of p53^4M^ ΔTAD and Random DNA during hollow condensate formation

For experimental conditions, a 10-μL reaction system was prepared by mixing 20 μM AF647-labeled p53^4M^ ΔTAD with 0.6 μM ATTO488-labeled Random DNA in a PCR tube, followed by incubation at RT for 30 minutes. Subsequently, 10 μL of droplet buffer containing p21 DNA (or 10 μL of droplet buffer without p21 DNA for the control group) was gently added to the system. This resulted in final concentrations of 10 μM p53^4M^ ΔTAD, 0.3 μM Random DNA, and 0.6 μM p21 DNA. The mixture was incubated at RT for a specified duration. After incubation, the reaction system was centrifuged at 13,800× g for 10 minutes at 20 °C. 5 μL supernatant was gently aspirated, diluted into 75 μL ddH₂O, and 75 μL mixture was transferred to a 384-well plate for fluorescence intensity measurements of AF647 and ATTO488 using a microplate reader (Varioskan LUX) (Supplementary Fig. 5a).

For the standard curves of AF647-labeled p53^4M^ ΔTAD and ATTO488-labeled Random DNA, concentration gradients of protein and DNA were separately prepared in 10 μL droplet buffer. After adding 10 μL of droplet buffer to each tube, the subsequent sample preparation and measurement were consistent with experimental conditions.

The microplate reader data were analyzed using Excel. Background subtraction was performed by subtracting the fluorescence results of the blank droplet buffer from all sample measurements. Standard curves for AF647-labeled p53^4M^ ΔTAD and ATTO488-labeled Random DNA were generated by plotting the known concentrations of protein or DNA against their corresponding fluorescence intensities, followed by linear regression analysis (Supplementary Fig. 5b). The fluorescence values of experimental samples were then interpolated into these standard curves to calculate the protein or DNA concentrations in the dilute phase of each experimental condition.

### Measurement of Young’s modulus of DPICs by AFM

DPIC samples were incubated at RT for 30 minutes after the addition of p21 DNA or Random DNA, followed by gentle rinsing with droplet buffer to remove loosely adhered DPICs from the coverslip surface. Measurements were performed with the operation mode of force volume mode in fluid on the commercial AFM BioScope Resolve (Bruker, Billerica, MA, USA), equipped with a silicon nitride probe (PFQNM-LC-A-CAL, Bruker; tip height: 17 μm, tip radius: 70 nm, spring constant: 0.083 N/m). Key parameters included a loading/unloading speed of 4 μm/s, scan size of 50 nm, 4 ramps per line, ramp size of 1 μm, ramp rate of 1 Hz, and a deflection error trigger threshold of 12 nm. 16 indentations were performed per DPIC. Experiments were completed at room temperature within 1 hour, with approximately 20 DPICs tested for each experiment.

Data were processed in Nanoscope Analysis software (Bruker), where three force curves (*n* = 3) with overlapping extend/retract baselines were selected per DPIC. After 0^th^ order baseline correction, force-time, height-time and separation-time curves were extracted, and Young’s modulus was calculated by fitting the extend curves to the Sneddon (Conical) contact model which was set in the software. This model was initially developed to calculate the Young’s modulus of a semi-infinite elastic plane ^20^, but has also been utilized in research to determine the Young’s modulus of cells ^21^. For our experimental system, the fitting results of this model reveal significant differences in the physical characteristics between the two types of DPICs.

### Determining the diffusion coefficient of Random DNA within and outside hollow DPIC by fluorescence correlation spectroscopy (FCS)

For sample preparation, AF647-labeled p53^4M^ ΔTAD was mixed with ATTO488-labeled Random DNA, followed by a 30-min incubation at room temperature on a coverslip surface. p21 DNA was subsequently introduced and subjected to 5 gently pipetting cycles to ensure maximum homogeneity of p21 DNA concentration. After a 120-min incubation, the prepared samples were then transferred to a commercial FCS setup (Olympus FV4000 confocal microscope with a CorTector CX-D20 FCS detection module). The fluorescent intensity-time curves of ATTO488-labeled Random DNA were recorded in the inner region (Lumen) and outer region (dilute phase) of hollow condensates. For each measurement point, 5 curves with 10-s duration were recorded.

The raw data was transformed to autocorrelation curves using Correlation Analysis V3.7 software. Relaxation times of autocorrelation curves were fitted by the model of Single 3D Diffusion with Triplet Dynamics, which enabled calculation of diffusion coefficients of ATTO488-labeled Random DNA.

### Unpaired t test

Statistical significance was evaluated based on Student’s t-tests (Prism 9 for macOS, Version 9.1.0 (216), March 15, 2021, GraphPad Software, Inc.). Test was chosen as unpaired, two-tailed t test. P value style: GP: 0.1234 (ns), 0.0332 (*), 0.0021 (**), 0.0002 (***), < 0.0001 (****).

### Box-plot

The function of “boxplot” in MATLAB software (R2015a, 64-bit, February 12, 2015) was used to plot the boxplots in Fig. 3, 4, and Supplementary Fig. 3. For each boxplot, the black line denotes the median, box edges represent the 25^th^ and 75^th^ percentiles, whiskers indicate the range excluding outliers, and outliers are shown as individual dots (•).

## References

1 Feric, M. & Misteli, T. Function moves biomolecular condensates in phase space. Bioessays 44, doi:10.1002/bies.202200001; ARTN e2200001 (2022).

2 Alberti, S. & Dormann, D. Liquid-Liquid Phase Separation in Disease. Annual Review of Genetics*, Vol* 53 53, 171–194, doi:10.1146/annurev-genet-112618-043527 (2019).

3 Bergmann, A. M. et al. Liquid spherical shells are a non-equilibrium steady state of active droplets. Nature Communications 14, 6552, doi:10.1038/s41467-023-42344-w (2023).

4 Alshareedah, I., Moosa, M. M., Raju, M., Potoyan, D. A. & Banerjee, P. R. Phase transition of RNA-protein complexes into ordered hollow condensates. P Natl Acad Sci USA 117, 15650–15658, doi:10.1073/pnas.1922365117 (2020).

5 Erkamp, N. A. et al. Spatially non-uniform condensates emerge from dynamically arrested phase separation. Nature Communications 14, 684, doi:10.1038/s41467-023-36059-1 (2023).

6 Li, C. et al. Deciphering the molecular mechanism underlying morphology transition in two-component DNA-protein cophase separation. Structure 33, 62–77.e68, doi:10.1016/j.str.2024.10.026 (2025).

7 Nikolova, P. V., Henckel, J., Lane, D. P. & Fersht, A. R. Semirational design of active tumor suppressor p53 DNA binding domain with enhanced stability. P Natl Acad Sci USA 95, 14675–14680, doi:DOI 10.1073/pnas.95.25.14675 (1998).

8 Wei, C. L. et al. A global map of p53 transcription-factor binding sites in the human genome. Cell 124, 207–219, doi:10.1016/j.cell.2005.10.043 (2006).

9 Hafner, A., Bulyk, M. L., Jambhekar, A. & Lahav, G. The multiple mechanisms that regulate p53 activity and cell fate. Nat Rev Mol Cell Bio 20, 199–210, doi:10.1038/s41580-019-0110-x (2019).

10 Cahn, J. W. & Hilliard, J. E. Free Energy of a Nonuniform System .1. Interfacial Free Energy. J Chem Phys 28, 258–267, doi:10.1063/1.1744102 (1958).

11 Lee, C. F., Brangwynne, C. P., Gharakhani, J., Hyman, A. A. & Jülicher, F. Spatial Organization of the Cell Cytoplasm by Position-Dependent Phase Separation (vol 111, 088101, 2013). Phys Rev Lett 111, doi:ARTN 269902; 10.1103/PhysRevLett.111.269902 (2013).

12 Hyman, A. A., Weber, C. A. & Juelicher, F. Liquid-Liquid Phase Separation in Biology. Annu Rev Cell Dev Bi 30, 39–58, doi:10.1146/annurev-cellbio-100913-013325 (2014).

13 Brangwynne, C. P., Tompa, P. & Pappu, R. V. Polymer physics of intracellular phase transitions. Nat Phys 11, 899–904, doi:10.1038/Nphys3532 (2015).

14 Berry, J., Brangwynne, C. P. & Haataja, M. Physical principles of intracellular organization via active and passive phase transitions. Reports on Progress in Physics 80, doi:ARTN 046601 10.1088/1361-6633/aaa61e (2018).

15 Sciortino, F., Bansil, R., Stanley, H. E. & Alstrom, P. Interference of Phase-Separation and Gelation - a Zeroth-Order Kinetic-Model. Phys Rev E 47, 4615–4618, doi:DOI 10.1103/PhysRevE.47.4615 (1993).

16 Shimobayashi, S. F., Ronceray, P., Sanders, D. W., Haataja, M. P. & Brangwynne, C. P. Nucleation landscape of biomolecular condensates. Nature 599, 503-+, doi:10.1038/s41586-021-03905-5 (2021).

17 Alberti, S., Gladfelter, A. & Mittag, T. Considerations and Challenges in Studying Liquid-Liquid Phase Separation and Biomolecular Condensates. Cell 176, 419–434, doi:10.1016/j.cell.2018.12.035 (2019).

18 Wen, P. et al. Coacervate vesicles assembled by liquid–liquid phase separation improve delivery of biopharmaceuticals. Nature Chemistry 17, 279–288, doi:10.1038/s41557-024-01705-8 (2025).

19 Liu, J. H., Zhorabek, F., Dai, X., Huang, J. Q. & Chau, Y. Minimalist Design of an Intrinsically Disordered Protein-Mimicking Scaffold for an Artificial Membraneless Organelle. Acs Central Sci 8, 493–500, doi:10.1021/acscentsci.1c01021 (2022).

20 Harding, J. W. & Sneddon, I. N. The elastic stresses produced by the indentation of the plane surface of a semi-infinite elastic solid by a rigid punch. Mathematical Proceedings of the Cambridge Philosophical Society 41, 16–26, doi:10.1017/S0305004100022325 (1945).

21 Sun, W. et al. Precise determination of elastic modulus of cell using conical AFM probe. J Biomech 118, 110277, 10.1016/j.jbiomech.2021.110277 (2021).

